# Amorphous nickel titanium alloy film: a new choice for cryo electron microscopy sample preparation

**DOI:** 10.1101/2020.03.02.963959

**Authors:** Xiaojun Huang, Lei Zhang, Zuoling Wen, Hui Chen, Shuoguo Li, Gang Ji, Chang-cheng Yin, Fei Sun

**Affiliations:** Center for Biological Imaging, Core Facilities for Protein Science, Institute of Biophysics, Chinese Academy of Sciences, Beijing 100101, China; Department of Biophysics, The Health Science Center, Peking University, Beijing 100191, China; Electron Microscopy Analysis Laboratory, Medical and Health Analysis Center, Peking University, Beijing 100191, China; National Key Laboratory of Biomacromolecules, CAS Center for Excellence in Biomacromolecules, Institute of Biophysics, Chinese Academy of Sciences, Beijing 100101, China; College of Life Science, University of Chinese Academy of Sciences, Beijing 100049, China; Center for Protein Science, Peking University, Beijing 100871, China; Key Laboratory of RNA Biology, CAS Center for Excellence in Biomacromolecules, Institute of Biophysics, Chinese Academy of Sciences, Beijing 100101, China

**Keywords:** Amorphous nickel titanium alloy, beam induced motion, cryoEM sample preparation, particle distribution, supporting film

## Abstract

Cryo-electron microscopy (cryoEM) has become one of the most important approach for structural biology. However, barriers are still there for an increase successful rate, a better resolution and improved efficiency from sample preparation, data collection to image processing. CryoEM sample preparation is one of the bottlenecks with many efforts made recently, including the optimization of supporting substrate (e.g. ultra-thin carbon, graphene, pure gold, 2d crystal of streptavidin, and affinity modification), which was aimed to solve air-water interface problem, or reduce beam induced motion (BIM), or change particle distribution in the grid hole. Here, we report another effort of developing a new supporting substrate, the amorphous nickel-titanium alloy (ANTA) film, for cryoEM sample preparation. Our investigations showed advantages of ANTA film in comparison with conventional carbon film, including superior electron conductivity and trace non-specific interaction with protein. These advantages yield less BIM and significantly improved particle distribution during cryoEM experiment of human apo-ferritn, thus resulting an improved reconstruction resolution from a reduced number of micrographs and particles. Unlike the pure gold film, the usage of the ANTA film is just same with the carbon film, compatible to conventional automatic cryoEM data collection procedure.

## INTRODUCTION

With breakthrough and developments in the past ten years, cryo-electron microscopy (cryoEM), especially cryoEM single particle analysis (SPA), has become a powerful technique to study the high resolution three dimensional structure of biological macromolecules in solution (Cheng et al., 2017). However, there are still many bottlenecks to overcome for the future wide application of this technique. One of the most important bottlenecks is how to routinely prepare suitable samples for high resolution data collection and the following structure determination (Glaeser, 2016).

During cryoEM sample preparation, we had many issues to challenge, including protein denature/degradation and preferred orientation induced by the air-water interface (Glaeser et al., 2016), non-uniform/ill distribution of protein particles in the hole (Drulyte et al., 2018), un-expected few number of particles in the hole (Snijder et al., 2017), not well controlled ice thickness and etc. These challenges increase difficulties and efficiencies for the subsequent cryoEM data collection and thus become the barrier of a better resolution. The way of cryoEM sample preparation also affects another important factor, beam induced motion (BIM), which is critical for high resolution cryoEM imaging (Brilot et al., 2012).

Many efforts have been made in the past to solve these challenges by improving cryoEM sample preparation. One approach is adjoining a layer of continuous film, e.g. ultra-thin carbon (Grassucci et al., 2007), graphene (Fan et al., 2019; Naydenova et al., 2019; Pantelic et al., 2012; Russo and Passmore, 2014a) or graphene derivate (Pantelic et al., 2010), and 2D crystal of streptavidin (Han et al., 2016; Wang et al., 2008) et al., on the top of the holey supporting film. This approach can solve air-water interface issue, alter particle distribution and improve BIM while additional experimental procedures are needed and the additional layer adds another noisy background of the final cryoEM image. Another approach is modifying the surface property of the holey supporting film, e.g. by glow discharging (Isabell et al., 1999), special treatment with surfactants or PEG (Meyerson et al., 2014) or antibody (Yu et al., 2016). This approach can improve the particle distribution in the hole. To overcome un-expected few number of particles in the hole, applying sample a number of times to saturate the interaction between the sample and the supporting film was successful (Snijder et al., 2017). In addition, besides the conventional carbon film, another attempt to change the supporting substrate to pure gold film was also successful for an extremely reduce BIM (Russo and Passmore, 2014b), while the polycrystalline property of the gold film adds another issue during routine cryoEM data collection. Besides difficulty of TEM alignment, the cryoEM image should be taken in the middle of the hole without the film edge when the pure gold film was used.

In this study, we developed a new type of metal supporting film, amorphous nickel-titanium alloy (ANTA) film, for another good choice of cryoEM sample preparation. The relevant product has been successfully used in real high resolution cryoEM applications (Hua et al., 2020; Wu et al., 2019). Our present studies further showed advantages of ANTA film in comparison to carbon film and pure gold film.

## MATERIALS AND METHODS

### Preparation of ANTA film and amorphous carbon film

The procedure of preparing supporting film with microarray holes was modified from the published method (Quispe et al., 2007). As shown in **Figure 1A**, the super elastic nickel titanium alloy wire of 0.5 mm diameter (TIANRUI NONFERROUS METAL, Beijing), which is consisted of 40 ∼ 60 atomic percent of nickel and 60 ∼ 40 atomic percent of titanium, was used to prepare the ANTA film. The wire was heated up to 1500 ∼ 2500 °C in a tungsten wire basket and evaporated to a silicon wafer with microarray holes. The evaporation was performed in an oil-free vacuum system (Beiyichuangxin, Beijing) with a high vacuum (< 5×10^−4^ Pa). The ANTA film was formed on the surface of silicon wafer.

**Figure 1.**
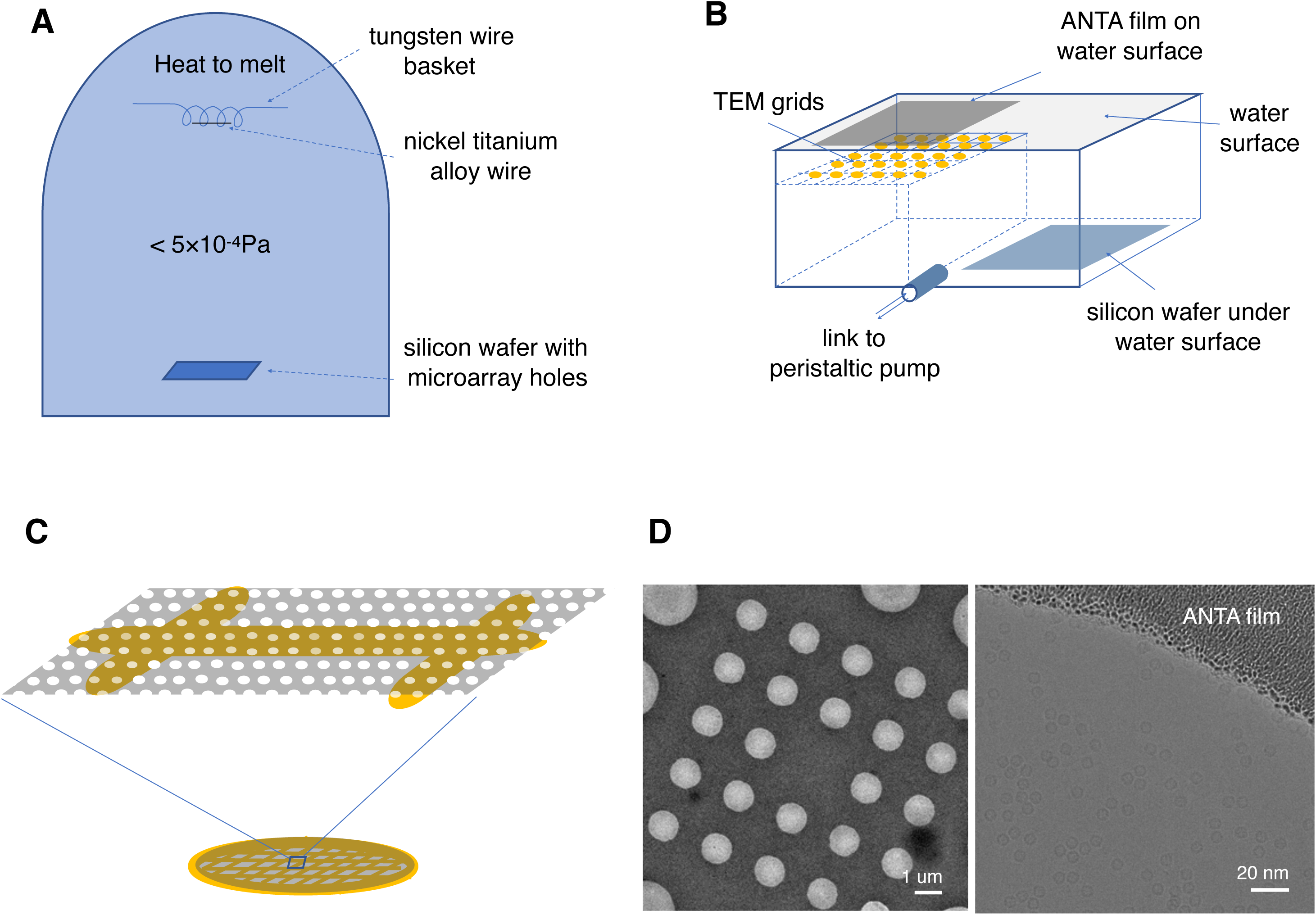
Preparation and usage of ANTA film. (**A**) Nickel titanium alloy wire is heated and evaporated under high vacuum to a fabricated silicon wafer with microarray holes. (**B**) The ANTA film on the silicon wafer is released on the water surface and transferred onto TEM grids (copper or gold). (**C**) A diagram to show how the ANTA film is attached onto one side of the TEM grid. The metal bar of the grid is colored in gold. (**D**) CryoEM images (low and high magnification) of ANTA film with human apo-ferritin vitrified on. The microarray hole pattern of ANTA film here is called R2-1 mix, which contains two kinds of hole sizes, 1 μm and 2 μm.

The preparation of amorphous C film was made by heating the graphite rod (Ted Pella, Lot#052514-17029) and evaporated at a similar condition to the preparation of ANTA film. The desired thickness of the supporting film was controlled between 18∼25 nm.

In order to release the supporting film, the silicon wafer was immersed into the pure water (**Figure 1B**). The air-water interface promotes the supporting film peel off and then float on the water surface. TEM grids were placed under the water surface and then coated with the supporting film by lowering the water level to allow grids attach to the film. Once the coated grids were taken out of water, they were dried and stored in cool and clean environment away from light (**Figure 1C**). The supporting film were examined using FEI Talos F200C transmission electron microscope (ThermoFisher Scientific) at 200kV with 4k*4k Ceta CMOS camera (**Figure 1D**).

### Film thickness measurement

Film thickness measurement was performed on a Nanoscope IIIa, MultiMode system SPM (Digital Instruments, Bruker) in a tapping mode and in air. Ultra-sharp NSC11/AIBS silicon cantilevers (MikroMasch, Tallinn, Estonia) with spring constants of ∼48 N/m and resonant frequencies of ∼330 kHz were used for imaging. Images were collected at a speed of 1 Hz with a resolution of 512×512 pixels. Two types of films were used for thickness measurement, the consecutive film and the microarray-hole one. The consecutive film was prepared by evaporating onto the freshly exposed surface of mica. To create a sharp edge for measurement, single sided Kapton Tape (Ted Pella, Prod No. 16090-6) was attached to the mica to cover the surface partially and removed after evaporation. The microarray-hole film was transferred to mica after peeling off from the silicon wafer. The existence of the holes allows the measurement at any place of the film. After imaging three random regions, flatten and section analysis were carried out using the Nanoscope analysis software to determine the thickness of the film.

### Electronic resistivity measurement

The nickel titanium alloy or amorphous carbon was evaporated onto the polished surface of a single-crystal silicon wafer. The thickness of the film was controlled as 30±3 nm and the size of the film was made as 1.8 cm × 1.8 cm. The resistance of the film was measured by four-wire resistance measurement using a physical property measurement system (Quantum Design, San Diego, CA). The distance between electrodes was kept 1 cm. The temperature variation was coupled with the resistance measurement from 300 K to 77 K and then back to 300 K. The resistivity was calculated according to the formula ρ=RS/L, where R is the measured electrical resistance, S is the cross-sectional area and L is the distance between the electrodes.

### Cryo sample preparation

TEM grids coated with ANTA film or carbon film were plasma cleaned in Gatan Solarus with the recipe of H_2_/O_2_ for 1min. 1.3 mg/ml human apo-ferritin (Hfn) in the buffer containing 50 mM Tris-HCl (pH8.0) and 150 mM NaCl was plunge frozen onto TEM grids using Leica EM GP. For the ANTA film, it is recommended to use a less force to touch the filter paper in comparison with carbon film, otherwise the ANTA film might peel off and attach to the filter paper.

Ryanodine receptor 1 (RyR1) was purified as the previous report (Wei et al., 2016) and stored in the buffer containing 20 mM HEPES (pH 7.2), 300 mM NaCl, 0.15 mg/ml amphipol and 2 mM DTT. Vitrification was performed using Vitrobot Mark IV (ThermoFisher Scientific) under 4 °C and 100% humidity. 3 μl of RyR1 at about 15 mg/ml was applied to ANTA copper 300-mesh grid or R1.2/1.3 Quantifoil 300-mesh copper grid, blotted for 5 s, and then plunge-frozen in liquid ethane.

### Measurement of beam induced motion (BIM)

BIM of each ANTA film and carbon film were measured using Hfn sample vitrified and the raw data were captured on FEI Titan Krios transmission electron microscope (ThermoFisher Scientific) at 300 kV with Falcon III camera (ThermoFisher Scientific) in pixel size of 1.42 Å. For each grid, 45 image stacks were taken at the centers of the corresponding holes from 3∼4 squares. Squares were selected from the center and four corner areas of the whole grid. Three grids of each group were used for the statistics. All image stacks were taken one minute after stage movement or stage tilt with the dose rate of 24 e/Å^2^/s, the exposure time of 1.1 s, the defocus range of −2 to −3 μm and the illumination area of ∼ 2um to cover the entire hole. The root of mean square (RMS) of the sample displacement was calculated according to the previous method (Zheng et al.,2017).

### CryoEM SPA data collection and processing

The cryoEM SPA datasets of Hfn sample were collected automatically using SerialEM (Mastronarde, 2005) on FEI Titan Krios transmission electron microscope (ThermoFisher Scientific) at 300 kV with Gatan K2 Summit camera. The pixel size was 0.41 Å in the super resolution mode, corresponding to the physical pixel size of 0.82 Å. Each image was exposed for 5.2 sec in a dose rate of 11.5 e/Å^2^/sec, resulting in a total dose of ∼ 60 e/ Å^2^. Each movie stack comprises 32 frames. The defocus range was set from −1 μm to −2.5 μm.

All movie stacks were motion-corrected, dose-weighted, distortion-corrected and binned to 0.82 Å by MontionCor2 (Zheng et al., 2017). CTF (Contrast Transfer Function) estimation was performed using CTFFIND4 (Rohou and Grigorieff, 2015). The subsequent data processing including particle picking, 2D and 3D classification, and 3D refinement was performed by using RELION 2.1 (Kimanius et al., 2016) and RELION 3.0 (Zivanov et al., 2018).

For the dataset using carbon film, 2,177 particles were manually picked from 43 micrographs and processed through reference-free 2D classification. Seven 2D class-averaged images were chosen as the references for automatic particle picking. In total, more than 90,000 particles were auto-picked from 747 micrographs. Then reference-free 2D classification and manual screening were carried out to remove overlapping or degraded particles. 64,667 particles were kept, extracted with a box size of 224 pixels and subjected into 3D auto-refinement, yielding a map with 3.2 Å resolution. After local CTF refinement, beam-tilt correction and particle re-extraction with box size of 288 pixels, the resolution was improved to 2.8 Å. The final map resolution was 2.6 Å after post processing. All reported resolutions were estimated based on the gold-standard FSC (Fourier Shell Correlation) at 0.143. All refinements were started from a 60 Å low-pass filtered initial model (PDB code, 2FHA).

For the dataset using ANTA film, the same workflow was applied. Briefly, 2,090 particles were initially manually picked from a subset and classified to create references for auto-picking. In total, more than 110,000 particles were auto-picked from 426 micrographs. After reference-free 2D classification and manual screening, 91,537 particles were selected for 3D refinement. The first round of 3D auto-refinement yielded a map of 3.0 Å resolution. After local CTF refinement and beam-tilt correction, the final resolution of the post processed map was improved to 2.4 Å.

### Model building and refinement

The atomic model of Hfm from the crystal structure (PDB code, 2FHA) was first fitted into the final cryoEM map as the initial model using Chimera (Pettersen et al., 2004). Then the real-space refinement was performed using PHENIX (Adams et al., 2010). The refined model was transformed into MRC files, and calculated FSC curves between the model and map by using software EMAN2 (Tang et al., 2007). The map visualization and segmentation were performed by software Chimera.

### Henderson-Rosenthal plot

To fairly compare cryoEM SPA results from the support of ANTA and carbon films, for each dataset, 45,211 particles were randomly selected by covering an even range of defocus and orientation distribution. Then partial (1%, 2%, 3%, 4%, 5%, 7%, 10%, 15%, 20%, 30%, 40%, 50%, 60%, 70%, 80% and 90%) datasets of total particles were selected. Reconstruction was then performed based on the specific subset of selected particles and the map resolution was estimated using the gold standard FSC_0.143_ criteria. Then the Henderson-Rosenthal plots for both datasets were generated as described (Stagg et al., 2014).

### Magnetic moment measurement of ANTA film

To measure the magnetic moment of ANTA film, the ANTA film was prepared on a single-crystal silicon wafer with the polished surface and the thickness of the film was controlled as 22 ± 3 nm and the size of the film was made as 1.8 cm × 1.8 cm. Then the magnetic moment was measured by SQUID (Superconducting Quantum Interference Device) using Magnetic Performance Measurement System (MPMS, Quantum Design Company) at the temperature of 77 K in the environmental magnetic strength of 0.0 T and 3.0 T, respectively. The magnetic moment of the silicon wafer in the same size was measured separately in the same condition as a control.

### Protein adhesion test with single molecule force spectroscopy

The carbon and ANTA were separately evaporated onto the clean cover slips (Fisherbrand Microscope cover glass, 12-545-C, 22×40-1, Fisher Scientific). The coating thickness was controlled as 30 nm. Single molecule force spectroscopy (SMFS) was performed using a home-built AFM as described previously (Lee, C. Y., et al., 2013). Biotinylated AFM cantilever (OBL-10, Bruker, Santa Barbara, CA) was ligated with the biotinylated human programmed death-ligand 1 (hPD-L1) by soluble formed streptavidin protein. The hPD-L1 ligated cantilever was provided by Dr. Jizhong Lou’s lab (Institute of Biophysics, CAS). Before each experiment, the hPD-L1 ligated cantilever was calibrated in PBS solution, showing a spring constant of around 4 pN/nm. The rupture force between hPD-L1 and the supporting film was measured by force ramp. In one measurement cycle, the AFM cantilever was brought into contact the supporting film for 200 ms, and then was retracted at 200 pN/s. The peak force was identified as the rupture force of the interaction between hPD-L1 and the supporting film.

### Hippocampal neuron cell culture on supporting films

The TEM grids coated with ANTA or carbon films were test for their ability of cell culture. The grids were placed in the culture dish with the side of the film up, sterilized under ultraviolet light overnight and coated with the Matrigel matrix (Product #354248, Corning, USA) just before usage.

The primary hippocampal neuron cells were separated from postnatal day 0 to day 1 Spraque-Dawley rats. The protocol of cell culture was modified from the published report (Kang et al., 2008). Hippocampal neurons were separated in HBSS buffer (H2387, Sigma) containing 20% FBS at 4 °C, then digested in the DNase/trypsin solution (0.5 mg/ml DNase, 5 mg/ml trypsin, 25 mM HEPES, 137 mM NaCl, 5 mM KCl, 7 mM Na_2_HPO_4_, pH 7.2) for 5 min at 37 °C. After the digestion, the single neuron cells were suspended in the DNase solution (0.5 mg/ml DNase, 25 mM HEPES, 137 mM NaCl, 5 mM KCl, 7 mM Na_2_HPO_4_, pH 7.2), and washed with HBSS containing 10% FBS.

After that, the primary single neuron cells were plated on the top of grids in the culture dish. The culture dish was kept at 37°C in 5% CO_2_ for one week. Culture media (DMEM supplemented with 10% fetal bovine serum, 1% penicillin and streptomycin) was changed every other day by refreshing half of the medium. The optical images of primary hippocampal neuron cells were acquired using an invert fluorescence microscope (Olympus IX81) with the 10X objective and sCMOS camera (Andor Zyla).

## RESULTS

### Thickness of ANTA film

The thickness of ANTA film is controlled by the distance between the source and the target as well as the length of the nickel titanium wire that will be depleted after the evaporation process. The exact thickness of film can be measured by atomic force microscope (Nanoscope System, South Korea) (**Figure 2**). Our measurements showed the thickness of ANTA film can be controlled between 18 to 25 nm when the distance from the source to the target is set to 9.0 cm and the length of the nickel titanium wire is 1.1 cm.

**Figure 2.**
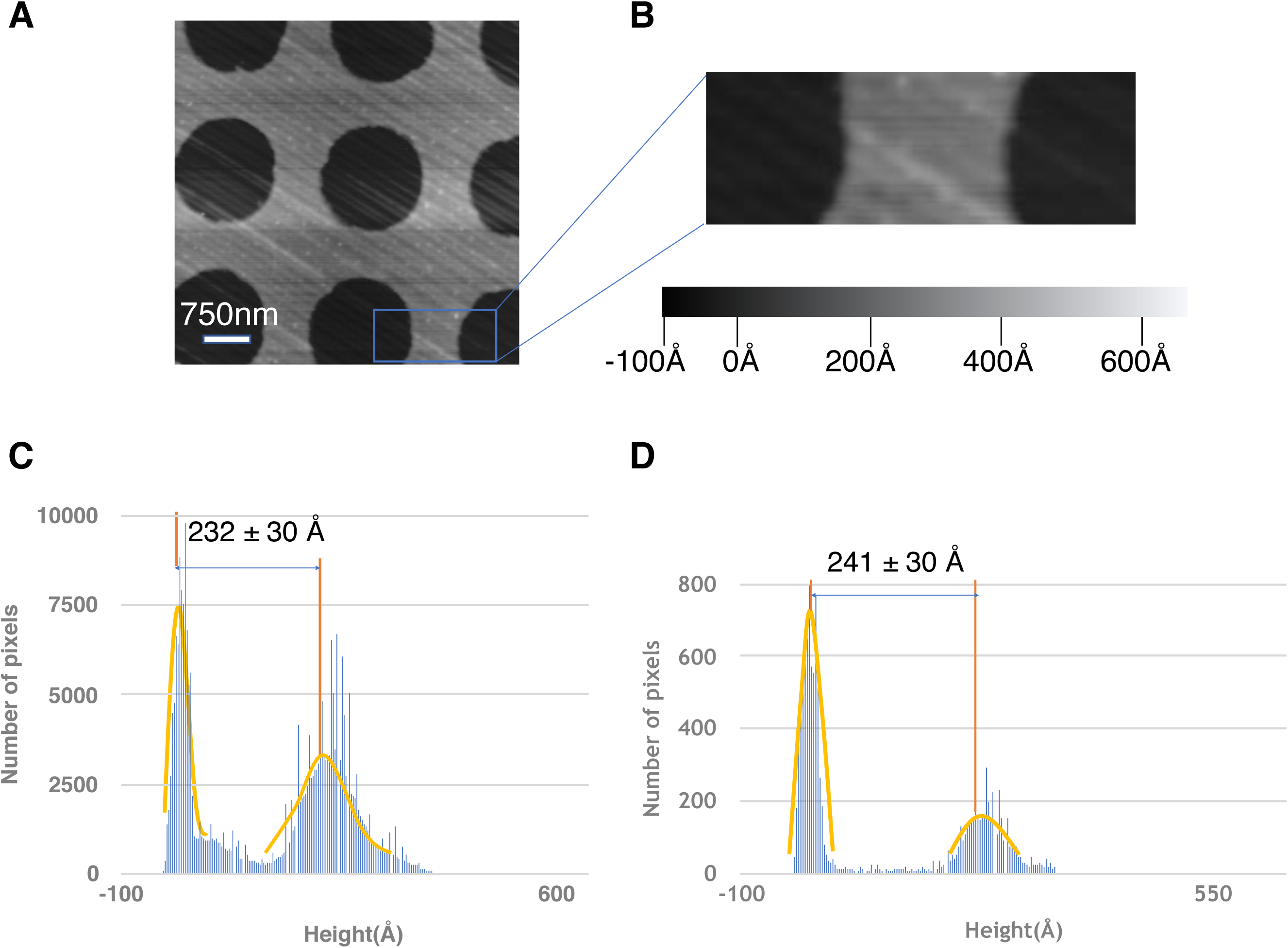
Thickness measurement of ANTA film by atomic force microscopy (AFM). (**A**) The topography image of ANTA film in a tapping-force mode. The film was transferred to a freshly exposed mica surface. (**B**) A zoom-in view of (A). The grey scale indicates the height. (**C**) The histogram of heights in (A). The two main peaks are fitted using Gaussian distribution. And the averaged thickness of ANTA film is measured as the distance between the two peaks. (**D**) The histogram of heights in (B).

### Low electrical resistivity of ANTA film at liquid nitrogen temperature

Compared to carbon film with the thickness of 30±3nm, the electrical resistivity of which increases significantly from 0.5×10^−2^Ω•m to 5×10^−2^Ω•m with the temperature decreases from 300 K to 77 K, the electrical resistivity of ANTA film is 4-order lower and does change significantly with the temperature, from 3.4×10^−6^Ω•m at 300 K to 3.8×10^−6^Ω•m at 77 K (**Figure 3A**). The superior electrical conductivity of ANTA film would yield a reduced beam induced motion (BIM) during cryoEM experiments in comparison with carbon film, which was proved from the dose dependent sample displacement measurements at both 0-degree stage tilt (**Figure 3B**) and 45-degree stage tilt (**Figure 3C**).

**Figure 3.**
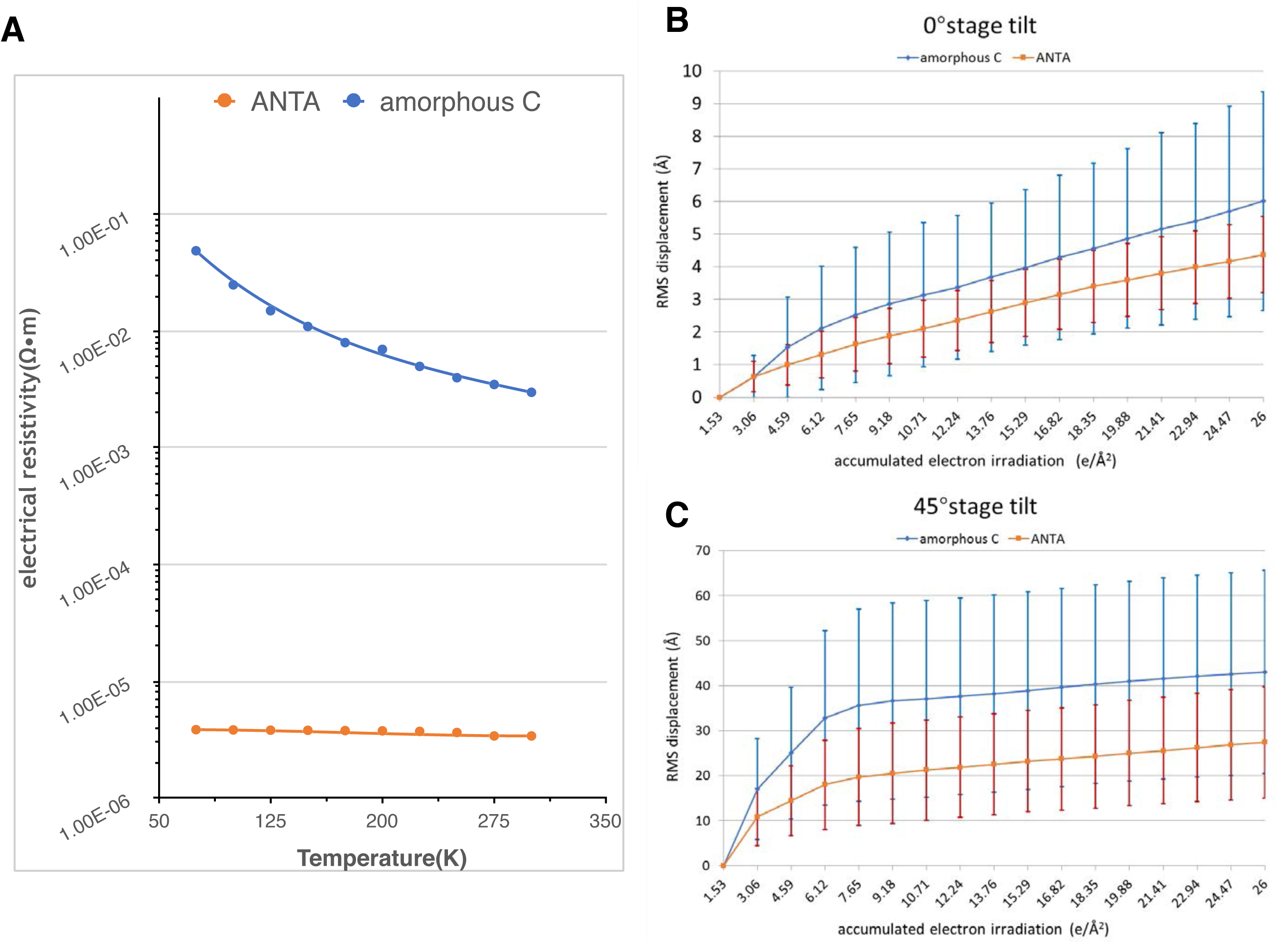
Electrical resistivity and beam induced motion (BIM) of ANTA film in comparison with carbon film. (A) Electrical resistivity of ANTA and carbon film under a range of temperature from 77 to 300 K. The thickness of each film is prepared as 30 nm. (**B**) and (**C**) The in-plane BIM of ANTA and carbon film under different electron irradiation dose. The experiments were performed under 90K with human apo-ferritin vitrified on the film. The stage was tilted zero degree in (B) and 45 degrees in (C), respectively. Each data point represents the average displacement of the whole imaging area captured and the error bar shows the standard deviation of all measurements (see Russo and Passmore, 2014). RMS, root of mean square.

### Trace magnetic moment of ANTA film

Because of containing the ferromagnetic element of nickel, it would be suspected that ANTA film would generate magnetic field that will affect electrons and thus influence cryoEM imaging significantly. Therefore, we measured the average magnetic moment of ANTA film on a silicon wafer. The experiments were performed in 77 K temperature to mimic the cryogenic condition. At the environmental magnetic strength of 0 T, the measured average magnetic moment in 148 s was −5.17×10^−6^ emu with standard deviation of 9.18×10^−9^ emu while at the environmental magnetic strength of 3 T, which mimics the environment inside objective lens of TEM, the measured average magnetic moment in 261 sec was 4.83×10^−4^ emu with standard deviation of 5.95×10^−7^ emu. As the control, the measured average magnetic moment of the silicon wafer in 242 sec at 0 T was 1.66×10^−9^ emu with the standard deviation of 7.93×10^−9^ emu, while at 3 T in 305 sec was −3.84×10^−5^ emu with the standard deviation of 4.61×10^−7^ emu. Our results showed that ANTA film has a trace magnetic moment, which will not yield obvious effects on cryoEM imaging.

### ANTA film is amorphous and suitable for routine TEM alignment

Previous studies have demonstrated the superior property of gold film with much less BIM and better for high resolution cryoEM SPA work. However, the polycrystalline property of gold film makes routine TEM alignment (including coma-free alignment and stigmatism adjustment) on gold film impossible (**Figure 4**), thus yielding additional difficulties for normal cryoEM SPA data collection. However, as another metal film, ANTA film is amorphous according to its electro diffraction pattern and the power spectrums of ANTA film images show the clear thon rings in response to the change of defocus values, which is just similar to carbon film (**Figure 4**). As a result, not like gold film, the ANTA film can be used in a same way with carbon film and suitable for defocus estimation, stigmatism adjustment and coma-free alignment.

**Figure 4.**
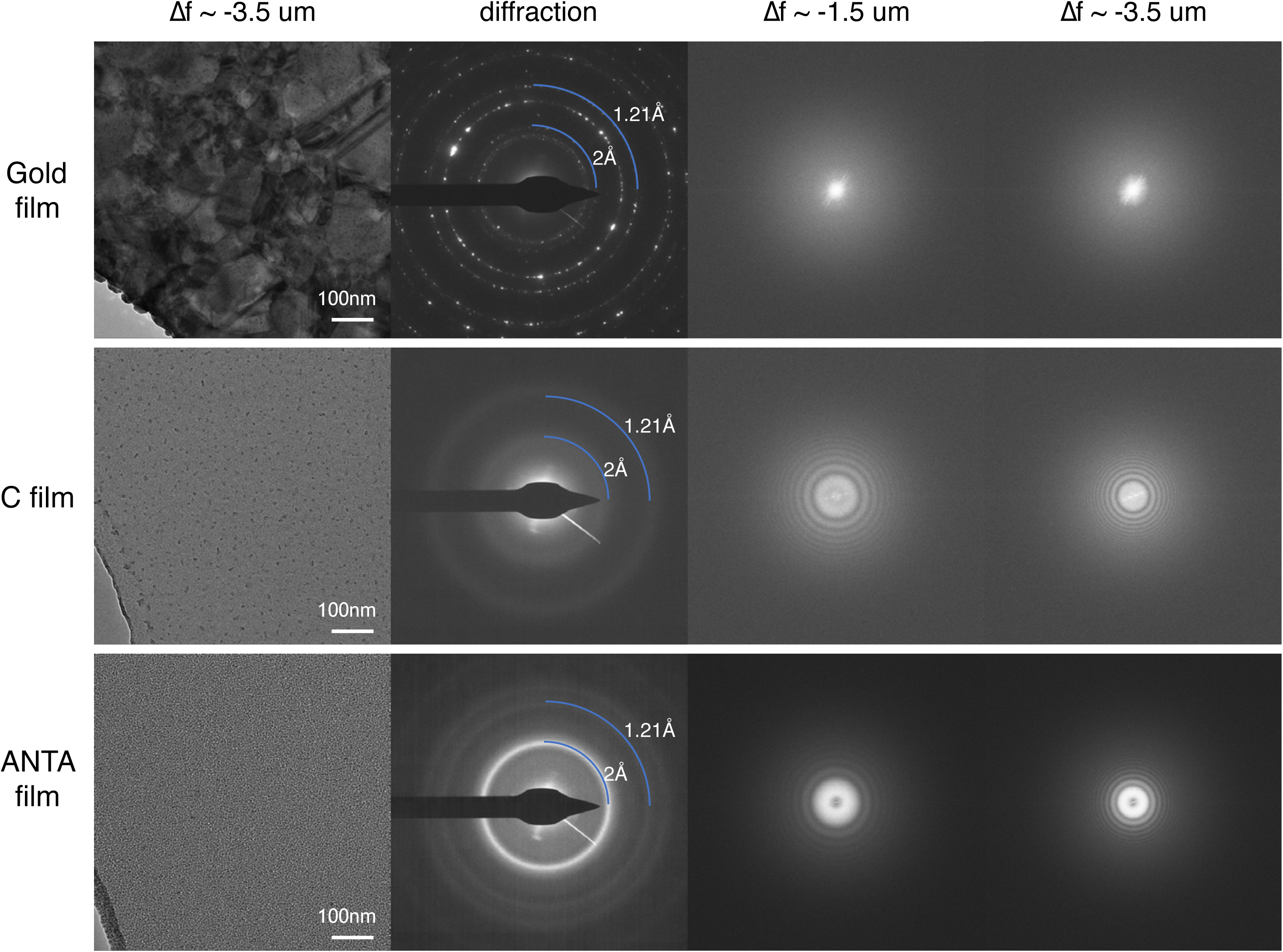
TEM characterization of ANTA film in comparison with polycrystalline gold and amorphous carbon films. The TEM micrographs of films (first column) were captured with the defocus of 3.5 μm and their power spectrums (third and fourth column) were measured with the defocus of 1.5 and 3.5 μm, respectively. The micrographs were taken at a magnification of 59,000x with a total dose of 50 e/ Å^2^. The electron diffraction patterns are shown in the second column. The electron diffraction images were taken at a camera length of 850 mm with beam stop inserted and a total dose of 15 e/ Å^2^.

### ANTA film yields better cryoEM SPA results in comparison with carbon film

Based on the reduced BIM of ANTA film in comparison with carbon film, we can predict cryoEM SPA experiments using ANTA film would yield a better result. Here, we selected Hfn as a test sample and collected cryoEM SPA datasets of the same batch of Hfn sample that were vitrified onto ANTA and carbon films, respectively. The final resolution of Hfn reaches 2.4 Å from 426 micrographs for ANTA film while that for carbon film reaches 2.6 Å from 747 micrographs (**Figures 5A, 5B and 5C**). To make a fair comparison, the same number of particles was selected from each dataset and the reconstruction was performed respectively. When the resolution for the carbon film dataset reaches 2.9 Å, the one for the ANTA film from the same number of particles reaches 2.7 Å (**Figure 5D**). The subsequent Henderson-Rosenthal analysis (**Figure 5E**) further confirmed that ANTA film can yield a better result using the same number of particles in comparison with carbon film.

**Figure 5.**
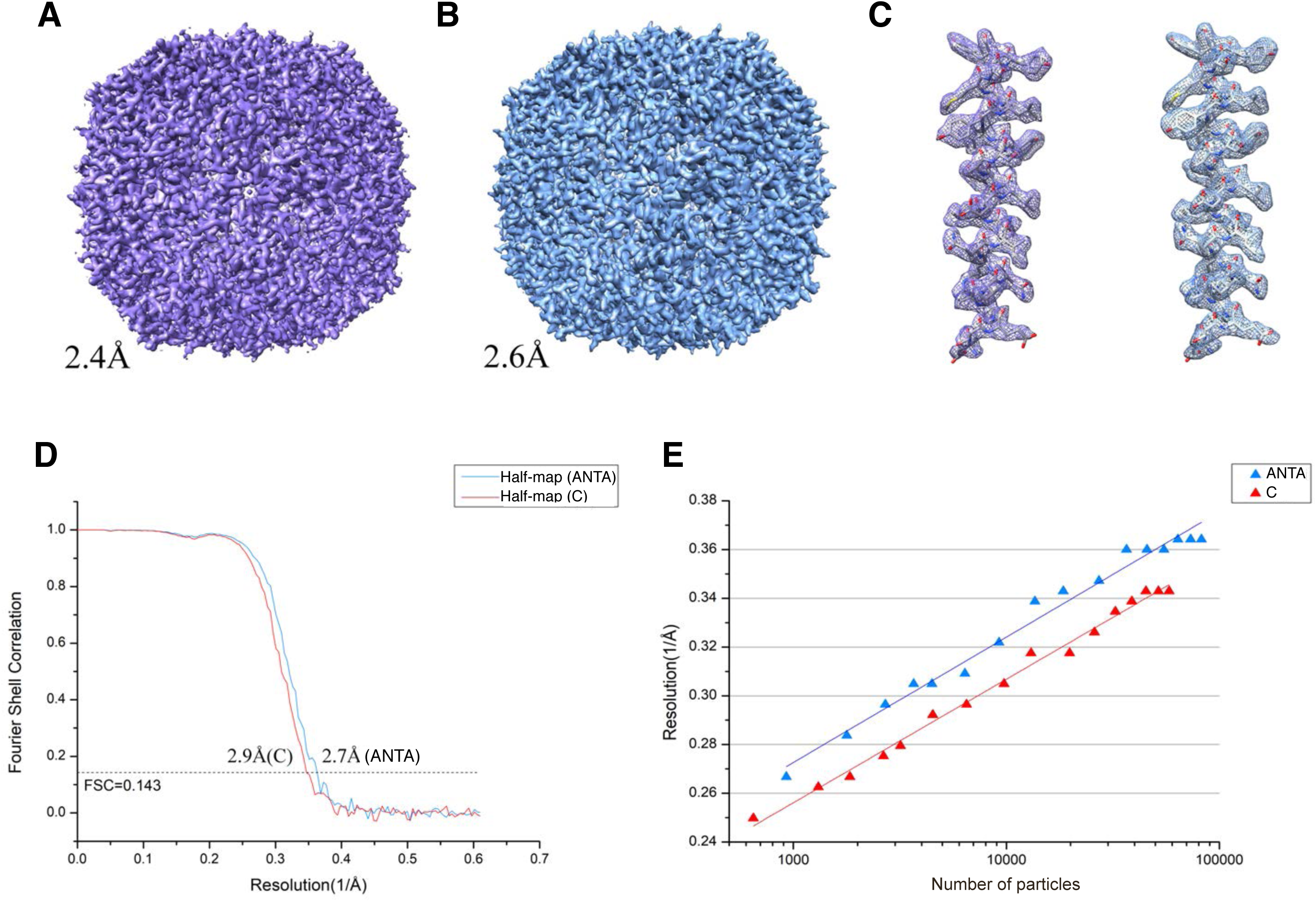
CryoEM SPA application of human apo-ferritin using ANTA film in comparison with carbon film. (**A**) and (**B**) CryoEM maps of human apo-ferritin reconstructed from the datasets collected using ANTA film grid (A) and carbon film grid (B), respectively. (**C**) Representative densities of both maps with atomic models fitted. Left, is from (A) and right from (B). (**D**) The gold-standard FSC (Fourier Shell Correlation) curves of both maps. (**E**) Henderson-Rosenthal plots for both cryoEM SPA experiments.

### Less none-specific protein absorption of ANTA film

We were interested to observe for ANTA film dataset more particles (∼110,000) can be picked from less (426) micrographs in comparison with carbon film dataset where ∼90,000 particles were picked from 747 micrographs. When we go back to investigate individual micrographs, we found the particle distribution in ANTA film dataset is more even and less aggregated in comparison with carbon film (**Figures 6A-6D**). We further selected another sample RyR1 to test this phenomenon and found (**Figures 6E and 6F**) there are more particles (∼ 100) consistently in the hole for the ANTA film grid while only a few (10 ∼ 30) can be picked for the carbon film grid (Quatifoil R1.2/1.3).

**Figure 6.**
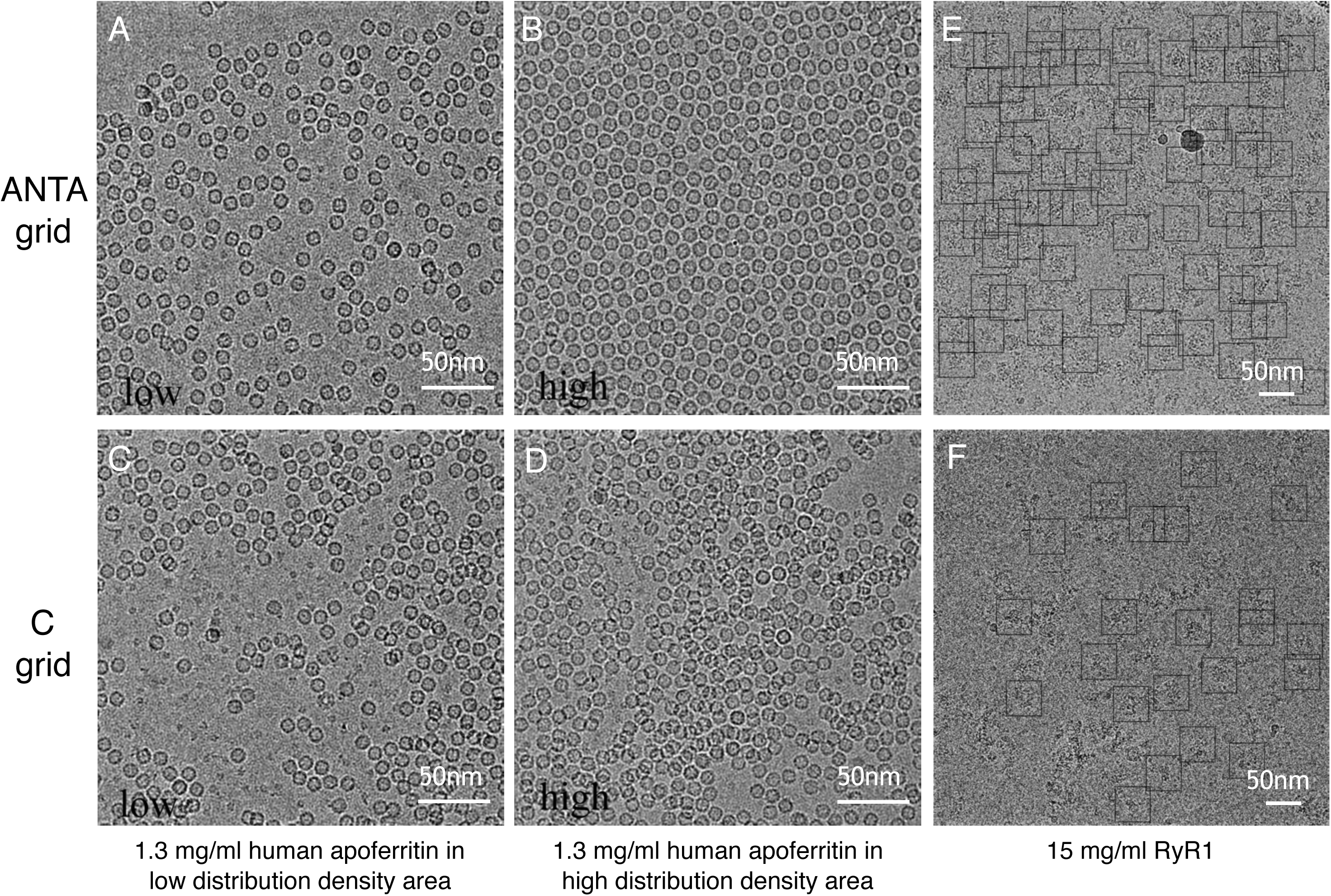
Particle distribution using ANTA film in comparison with carbon film. (**A**) and (**B**) CryoEM micrographs of 1.3 mg/ml human apo-ferrtin frozen on the same ANTA film grid were taken at both low (A) and high (B) distribution density areas. (**C**) and (**D**) CryoEM micrographs of 1.3 mg/ml human apoferrtin frozen on the same carbon film grid were taken at both low (C) and high (D) distribution density areas. (**E**) and (**F**) Typical cryoEM micrographs of 15 mg/ml ryanodine receptor type 1 (RyR1) vitrified using ANTA film grid (E) and carbon grid (F). More than 100 particles can be picked from (E) while less than 30 from (F).

We thought this phenomenon might be due to a less none-specific protein absorption of ANTA film in comparison with carbon film. For carbon film, more protein particles are presumably absorbed by carbon film and such absorption would also induce aggregation of the particles. For prove this concept, we performed the single molecule force spectroscopy to measure the interaction force between protein and the supporting film. In the current experiment (**Figure 7**), we selected human programmed death-ligand 1 (hPD-L1) as one testing example. The results showed that the averaged dissociation force was 16.6 pN between hPD-L1and the carbon film while the force between hPD-L1and ANTA film was only 0.94 pN (**Figure 7**).

**Figure 7.**
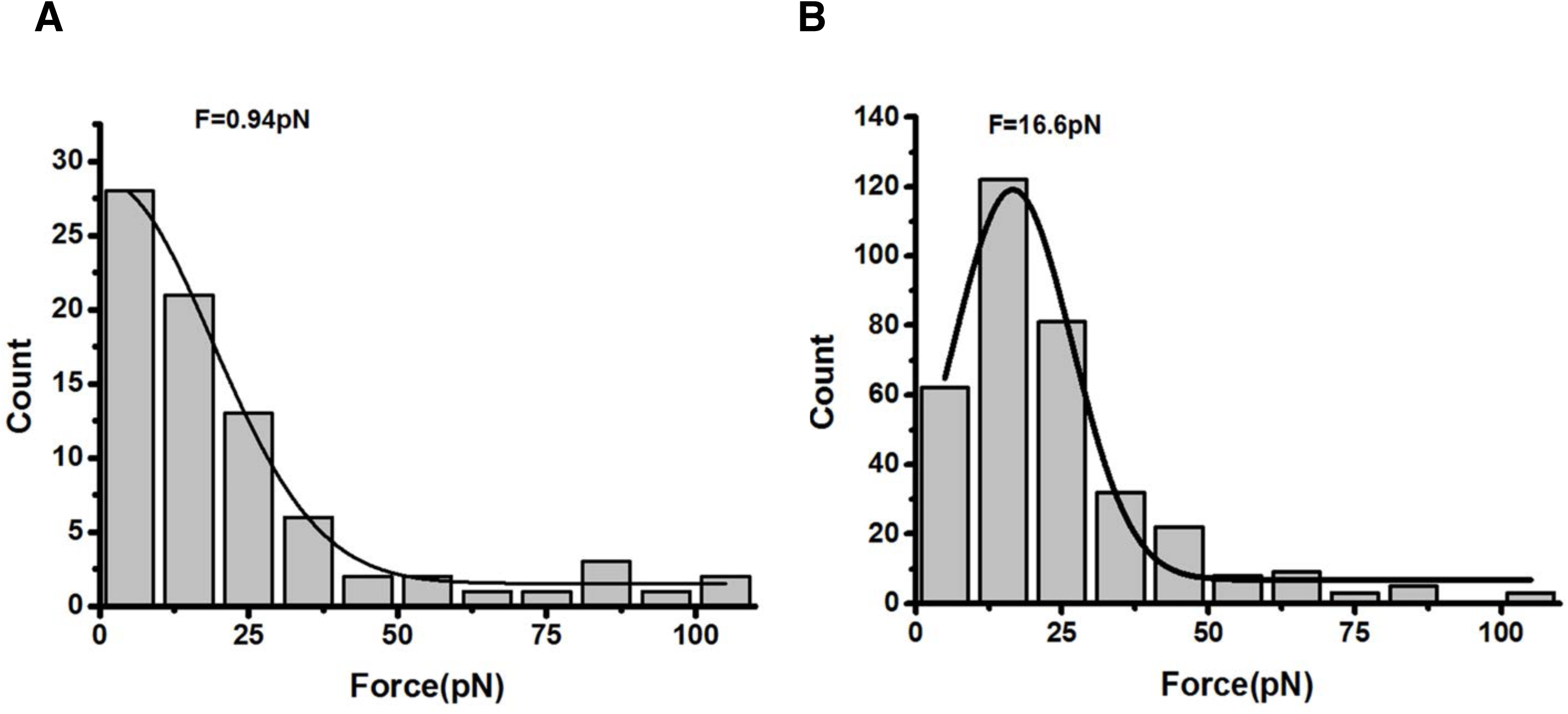
None-specific interaction between ANTA film and protein in comparison with carbon film. The statistics of rupture force interaction between HPD-L1 and the supporting film was measured using AFM. The histograms of force measurements are shown for ANTA film (A) and carbon film (B), respectively. A Gaussian fit was performed to calculated the averaged none-specific interaction force between HPD-L1 and the supporting film.

### Good biocompatibility of ANTA film

The above experiments have showed that ANTA film is suitable for cryoEM SPA experiments and better than carbon film. We ought to investigate the possibility of ANTA film for the application of *in situ* cryo-electron tomography and whether it is toxic for cell culturing. We tried to culture rat hippocampal neuron cells, very sensitive to toxic environment, onto the gold grids coated with ANTA and carbon films, respectively. The results showed that the cells can grow and differentiate well on both films (**Figure 8**), demonstrating a similar biocompatibility of ANTA film to carbon film.

**Figure 8.**
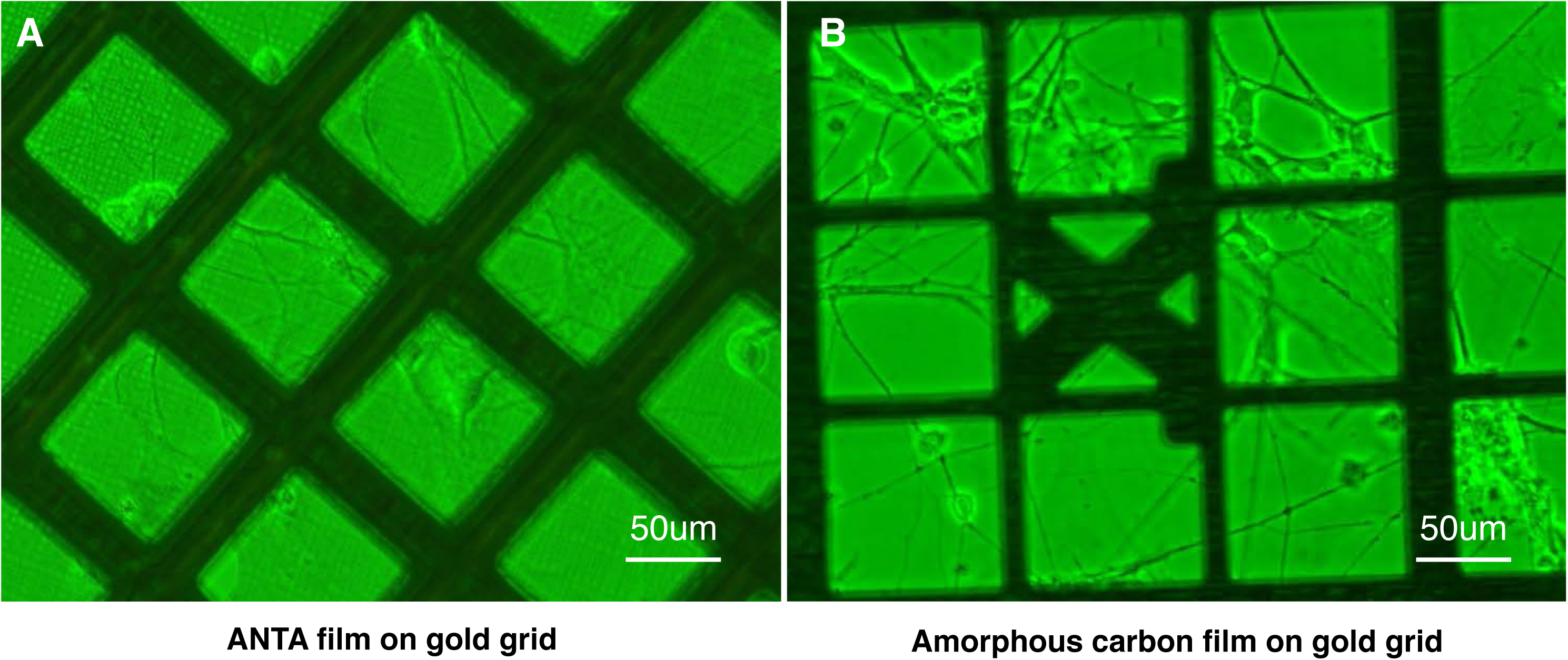
Culturing hippocampal neuron cell on both ANTA film (A) and carbon film (B). Both films are coated onto gold grids.

## DISCUSSIONS

In the present study, we developed a new supporting film for cryoEM experiments, the amorphous nickel-titanium alloy film (ANTA film). As a metal supporting film, the electrical conductivity of ANTA film is much greater than carbon film especially at liquid nitrogen temperature. Therefore, ANTA film exhibits less BIM than carbon film. As a result, in cryoEM SPA application using the support of ANTA film, we could achieve better reconstruction resolution from the same number of particles in comparison with the support of carbon film.

Compared to the pure gold film, another currently widely used metal supporting film, ANTA film is amorphous not polycrystalline, which enables us to perform routine TEM alignment (focus estimation, stigmatism correction and coma-free alignment) directly on the supporting film in a similar way to carbon film, and compatible to the current automatic cryoEM SPA data collection strategy.

However, according to the previous research (Russo and Passmore, 2014), the BIM effect using ANTA film is still a bit higher than the pure gold film, which would be explained by two reasons. First, the electrical conductivity of gold is better than ANTA, about two orders of magnitude lower in the electrical resistivity. Second, ANTA film is coated onto a gold/copper grid and the slightly different thermal expansion coefficient between the supporting film and the grid would induce mechanical instability after vitrification while the pure gold grid does not have such problem.

The thermal expansion coefficient of ANTA film is expected to be between nickel (13.4×10^−6^ K^-1^) and titanium (8.6×10^−6^ K^-1^). The thermal expansion coefficient of block nickel titanium alloy with a similar atomic ratio is around 11×10^−6^ K^-1^. To reduce cryo-crinkling during vitrification, the selected material of grid should have a similar thermal expansion coefficient with the ANTA film. The grid made of stainless steel (11.8 × 10^−6^ K^-1^) is most suitable for ANTA film coating. The grids made of nickel and titanium are also acceptable. In general, the gold grid with the thermal expansion coefficient of 14.2×10-6 K-1 can also be used and it is better than the copper grid with the thermal expansion coefficient of 16.5×10-6 K-1. To be noted, the above thermal expansion coefficients are looked up from WebElements (http://www.webelements.com/).

The high contrast of ANTA supporting film during cryoEM experiments makes identification of the hole area much easier (**Figure 1D**), which is because the atomic numbers of both nickel and titanium are higher than the elements of biological sample. However, it is harder to distinguish the thickness of ice in the hole by comparing the contrast of the hole area with the film area. The similar effect can be also observed when using the pure gold grid.

The most interesting finding of the current study is the surface property of ANTA film that is significantly different from carbon film. Using hPD-L1 as a test sample, the single molecule force spectroscopy showed a strong none-specific interaction between carbon film and the hPD-L1 protein. The measured force (16.6 pN in average) is strong enough to absorb protein particles and induce protein denature in a high probability, which might explain the non-uniform distribution and aggregation effect of Hfn particles observed from cryoEM micrographs. However, the measured force (0.94 pN in average) between ANTA film and the hPD-L1 protein is less than 1 pN, which is very small and would not induce significant change of the interacted particles. The strong non-specific interaction between carbon film and protein sample would be due to the hydrophobic nature of carbon film, which is easy to form hydrophobic interaction with the hydrophobic surface of protein sample and also dangerous for protein denature. While, as a metal film, the surface of ANTA film is hydrophilic, which is favorable to reduce none-specific interaction with protein sample. In our many applications (RyR1 sample in the present study), we found using carbon film it is rare to allow the particles distribute in the hole while the case can be efficiently solved by replacing to ANTA film. This observation would be due to the different surface property between ANTA film and carbon film, yielding another advantage of ANTA film.

It would be susceptible for the toxicity of ANTA film because of containing nickel atoms. However, our experiments showed a good biocompatibility of ANTA film suitable for cell growth. This observation is due to the formation of titanium oxide layer on the alloy surface (Figueira et al., 2009), which blocks the release of nickel ion and eliminates the potential negative effects on cell culture, yielding a good corrosion resistance of ANTA film.

The potential ferromagnetic property of ANTA film is another concern for the application of ANTA film. However, our measurement of magnetic moment of ANTA film is much small compared to the strong magnetic field in the objective lens and thus can be ignored. In addition, our cryoEM SPA experiments showed we can achieve 2.4 Å resolution for Hfn sample using ANTA film. As a result, in the resolution range of 2∼3 Å, the potential magnetic effect from ANTA film is not necessary to consider.

Overall, ANTA film provides another choice for cryoEM sample preparation and has the potential to improve the sample distribution and increase the efficiency of data collection in comparison with carbon film.

## DATA DEPOSITION

The final cryoEM maps of human apo-ferritin from datasets using the grid support of ANTA film and carbon film have been deposited into Electron Microscope Data Bank (EMDB) with the accession codes EMD-XXXX and EMD-XXXX, respectively. The corresponding datasets have been deposited into EMPIAR with the accession codes EMPIAR-XXXX and EMPIAR-XXXX, respectively.

## ACKNOWLEDMENTS

We would like to thank Kelong Fan and Minmin Liang from Dr. Xiyun Yan’s lab at Institute of Biophysics, CAS for preparing high-quality apoferritin sample, Dr. Wei Ding from Institute of physics, CAS for preliminary data analysis and Dr. Jizhong Lou from Institute of Biophysics, CAS for advices of single molecule force spectroscopy experiment. All the sample preparation and cryoEM works were performed at Center for Biological Imaging (CBI, http://cbi.ibp.ac.cn), Institute of Biophysics, Chinese Academy of Sciences.

This work was supported by grants from National Natural Science Foundation of China (31830020 and 31925026 to FS, and 31500608 to XH), the Ministry of Science and Technology of China (2017YFA0504702) to GJ, Beijing Municipal Science and Technology Commission (Z181100004218002) to FS, and Technological Innovation Program of Chinese Academy of Sciences (Y5CZ023001) to XH.

## REFERENCES

Adams, P.D., Afonine, P.V., Bunkoczi, G., Chen, V.B., Davis, I.W., Echols, N., Headd, J.J., Hung, L.W., Kapral, G.J., Grosse-Kunstleve, R.W., McCoy, A.J., Moriarty, N.W., Oeffner, R., Read, R.J., Richardson, D.C., Richardson, J.S., Terwilliger, T.C. and Zwart, P.H., 2010. PHENIX: a comprehensive Python-based system for macromolecular structure solution, Acta Crystallogr D Biol Crystallogr. 66, 213–21.

Brilot, A.F., Chen, J.Z., Cheng, A., Pan, J., Harrison, S.C., Potter, C.S., Carragher, B., Henderson, R. and Grigorieff, N., 2012. Beam-induced motion of vitrified specimen on holey carbon film, J Struct Biol. 177, 630–7.

Cheng, Y., Glaeser, R.M. and Nogales, E., 2017. How Cryo-EM Became so Hot, Cell. 171, 1229–1231.

Drulyte, I., Johnson, R.M., Hesketh, E.L., Hurdiss, D.L., Scarff, C.A., Porav, S.A., Ranson, N.A., Muench, S.P. and Thompson, R.F., 2018. Approaches to altering particle distributions in cryo-electron microscopy sample preparation, Acta Crystallogr D Struct Biol. 74, 560–571.

Fan, X., Wang, J., Zhang, X., Yang, Z., Zhang, J.C., Zhao, L., Peng, H.L., Lei, J. and Wang, H.W., 2019. Single particle cryo-EM reconstruction of 52 kDa streptavidin at 3.2 Angstrom resolution, Nat Commun. 10, 2386.

Figueira, N., Silva, T.M., Carmezim, M.J. and Fernandes, J.C.S., 2009. Corrosion behaviour of NiTi alloy, Electrochimica Acta. 54, 921–926.

Glaeser, R.M., 2016. How good can cryo-EM become?, Nat Methods. 13, 28–32.

Glaeser, R.M., Han, B.G., Csencsits, R., Killilea, A., Pulk, A. and Cate, J.H., 2016. Factors that Influence the Formation and Stability of Thin, Cryo-EM Specimens, Biophys J. 110, 749–55.

Grassucci, R.A., Taylor, D.J. and Frank, J., 2007. Preparation of macromolecular complexes for cryo-electron microscopy, Nat Protoc. 2, 3239–46.

Han, B.G., Watson, Z., Kang, H., Pulk, A., Downing, K.H., Cate, J. and Glaeser, R.M., 2016. Long shelf-life streptavidin support-films suitable for electron microscopy of biological macromolecules, Journal of Structural Biology. 195, 238–244.

Hua, T., Li, X., Wu, L., Iliopoulos-Tsoutsouvas, C., Wang, Y., Wu, M., Shen, L., Johnston, C.A., Nikas, S.P., Song, F., Song, X., Yuan, S., Sun, Q., Wu, Y., Jiang, S., Grim, T.W., Benchama, O., Stahl, E.L., Zvonok, N., Zhao, S., Bohn, L.M., Makriyannis, A. and Liu, Z.J., 2020. Activation and Signaling Mechanism Revealed by Cannabinoid Receptor-Gi Complex Structures, Cell. 180, 655–665 e18.

Isabell, T.C., Fischione, P.E., O’Keefe, C., Guruz, M.U. and Dravid, V.P., 1999. Plasma Cleaning and Its Applications for Electron Microscopy, Microsc Microanal. 5, 126–135.

Kang, J.S., Tian, J.H., Pan, P.Y., Zald, P., Li, C., Deng, C. and Sheng, Z.H., 2008. Docking of axonal mitochondria by syntaphilin controls their mobility and affects short-term facilitation, Cell. 132, 137–48.

Kimanius, D., Forsberg, B.O., Scheres, S.H. and Lindahl, E., 2016. Accelerated cryo-EM structure determination with parallelisation using GPUs in RELION-2, Elife. 5.

Mastronarde, D.N., 2005. Automated electron microscope tomography using robust prediction of specimen movements, J Struct Biol. 152, 36–51.

Meyerson, J.R., Rao, P., Kumar, J., Chittori, S., Banerjee, S., Pierson, J., Mayer, M.L. and Subramaniam, S., 2014. Self-assembled monolayers improve protein distribution on holey carbon cryo-EM supports, Sci Rep. 4, 7084.

Naydenova, K., Peet, M.J. and Russo, C.J., 2019. Multifunctional graphene supports for electron cryomicroscopy, Proc Natl Acad Sci U S A. 116, 11718–11724.

Pantelic, R.S., Meyer, J.C., Kaiser, U., Baumeister, W. and Plitzko, J.M., 2010. Graphene oxide: A substrate for optimizing preparations of frozen-hydrated samples, Journal of Structural Biology. 170, 152–156.

Pantelic, R.S., Meyer, J.C., Kaiser, U. and Stahlberg, H., 2012. The application of graphene as a sample support in transmission electron microscopy, Solid State Communications. 152, 1375–1382.

Pettersen, E.F., Goddard, T.D., Huang, C.C., Couch, G.S., Greenblatt, D.M., Meng, E.C. and Ferrin, T.E., 2004. UCSF Chimera--a visualization system for exploratory research and analysis, J Comput Chem. 25, 1605–12.

Quispe, J., Damiano, J., Mick, S.E., Nackashi, D.P., Fellmann, D., Ajero, T.G., Carragher, B. and Potter, C.S., 2007. An improved holey carbon film for cryo-electron microscopy, Microscopy and Microanalysis. 13, 365–371.

Rohou, A. and Grigorieff, N., 2015. CTFFIND4: Fast and accurate defocus estimation from electron micrographs, J Struct Biol. 192, 216–21.

Russo, C.J. and Passmore, L.A., 2014a. Controlling protein adsorption on graphene for cryo-EM using low-energy hydrogen plasmas, Nat Methods. 11, 649–52.

Russo, C.J. and Passmore, L.A., 2014b. Electron microscopy: Ultrastable gold substrates for electron cryomicroscopy, Science. 346, 1377–80.

Snijder, J., Borst, A.J., Dosey, A., Walls, A.C., Burrell, A., Reddy, V.S., Kollman, J.M. and Veesler, D., 2017. Vitrification after multiple rounds of sample application and blotting improves particle density on cryo-electron microscopy grids, J Struct Biol. 198, 38–42.

Stagg, S.M., Noble, A.J., Spilman, M. and Chapman, M.S., 2014. ResLog plots as an empirical metric of the quality of cryo-EM reconstructions, J Struct Biol. 185, 418–26.

Tang, G., Peng, L., Baldwin, P.R., Mann, D.S., Jiang, W., Rees, I. and Ludtke, S.J., 2007. EMAN2: an extensible image processing suite for electron microscopy, J Struct Biol. 157, 38–46.

Wang, L.G., Ounjai, P. and Sigworth, F.J., 2008. Streptavidin crystals as nanostructured supports and image-calibration references for cryo-EM data collection, Journal of Structural Biology. 164, 190–198.

Wu, C., Huang, X., Cheng, J., Zhu, D. and Zhang, X., 2019. High-quality, high-throughput cryo-electron microscopy data collection via beam tilt and astigmatism-free beam-image shift, J Struct Biol. 208.

Yu, G., Li, K. and Jiang, W., 2016. Antibody-based affinity cryo-EM grid, Methods.

Zheng, S.Q., Palovcak, E., Armache, J.P., Verba, K.A., Cheng, Y. and Agard, D.A., 2017. MotionCor2: anisotropic correction of beam-induced motion for improved cryo-electron microscopy, Nat Methods. 14, 331–332.

Zivanov, J., Nakane, T., Forsberg, B.O., Kimanius, D., Hagen, W.J., Lindahl, E. and Scheres, S.H., 2018. New tools for automated high-resolution cryo-EM structure determination in RELION-3, Elife. 7.

